# Origin Matters: Using a Local Reference Genome Improves Measures in Population Genomics

**DOI:** 10.1101/2023.01.10.523267

**Authors:** Doko-Miles J. Thorburn, Kostas Sagonas, Mahesh Binzer-Panchal, Frederic J.J. Chain, Philine G.D. Feulner, Erich Bornberg-Bauer, Thorsten BH Reusch, Irene E. Samonte-Padilla, Manfred Milinski, Tobias L. Lenz, Christophe Eizaguirre

## Abstract

Genome-level sequencing enables us to ask fundamental questions about the genetic basis of adaptation, population structure, and epigenetic mechanisms, but usually requires a suitable reference genome for mapping population-level re-sequencing data. In some model systems, multiple reference genomes are available, giving researchers the challenging task of determining which reference genome best suits their data. Here we compare the use of two different reference genomes for the three-spined stickleback (*Gasterosteus aculeatus*), one novel genome derived from a European gynogenetic individual and the published reference genome of a North American individual. Specifically, we investigate the impact of using a local reference versus one generated from a distinct lineage on several common population genomics analyses. Through mapping genome resequencing data of 60 sticklebacks from across Europe and North America, we demonstrate that genetic distance among samples and the reference impacts downstream analyses. Using a local reference genome increased mapping efficiency and genotyping accuracy, effectively retaining more and better data. Despite comparable distributions of the metrics generated across the genome using SNP data (i.e., π, Tajima’s *D*, and *F*_ST_), window-based statistics using different references resulted in different outlier genes and enriched gene functions. A marker-based analysis of DNA methylation distributions had a comparably high overlap in outlier genes and functions, yet with distinct differences depending on the reference genome. Overall, our results highlight how using a local reference genome decreases reference bias to increase confidence in downstream analyses of the data. Such results have significant implications in all reference-genome-based population genomic analyses.

## Introduction

Genome-level sequencing has revolutionized many biological fields including evolution, ecology, microbiology, and population genomics (M. R. Jones & Good, 2016; Kao et al., 2014; Stapley et al., 2010). Historically, scientists have relied on one or a small number of high-quality reference genomes to address their specific questions. Progressively, the availability of high-quality reference genomes from large-scale projects (e.g., Earth BioGenome and Vertebrate Genome Projects; Lewin et al., 2018; Scientists, 2009), the decreasing costs of sequencing, and the availability of curated variant databases (e.g., dbSNP and dbVar; Lappalainen et al., 2013; Sherry et al., 2001) have improved the breadth and depth of genomic research. However, limitations due to the availability and quality of these resources and the challenge of integrating and interpreting multiple sources of genetic variation still exist (Formenti et al., 2022). These problems are only exacerbated in non-model systems where a high-quality reference genome may not exist. Recently, a graph-based genome alignment methodology has been developed to index and incorporate variant databases, providing a practical and highly efficient method that better captures variation in a genome (Kim et al., 2019). However, databases of high-quality variants are not currently available for most study systems. Moreover, in highly variable regions graph-based approaches can incur significant computational overhead and have an increased change of false-positive alignments (Grytten et al., 2020; Pritt et al., 2018). Hence, until graph-based methods become common place, assembling a haploid reference is still a viable alternative to making and utilising a resource intensive genome graph.

Reference genomes are integral parts of the analytical frameworks of genomic research. Specifically, variants are identified and referred to by their loci in relation to their position mapped on a reference genome. Such an approach allows for direct comparisons among multiple individuals and organisms, therein enabling comprehensive research in various fields, including phylogenomics (Fang et al., 2018; Wang et al., 2013), comparative evolution (Palmer & Kronforst, 2020; Parker et al., 2013), adaptive radiations (Ronco et al., 2020), and the genomics of domestication (Frantz et al., 2015).

Arguably one of the most important factors to consider when multiple reference genomes or assembly versions are available is their difference in quality. Whilst we are moving towards complete and error-free assemblies (Rhie et al., 2021), the continuing advances of methodologies can create significant differences among assemblies. For example, bioinformatic tools have been developed to resolve false gene duplications that stem from heterozygosity in homologous haplotypes (Guan et al., 2020; Roach et al., 2018). In some cases, such as in humans, reference genomes are continually updated alongside major advances, where choosing the most updated version will offer the most accurate analysis of human sequencing data (Guo et al., 2017; International Human Genome Sequencing Consortium, 2001, 2004; Pushkarev et al., 2009). However, when multiple similar quality and genetically diverse reference genomes are available from multiple populations or strains (e.g., Berner et al., 2019; Gan et al., 2011; Hirsch et al., 2016; Springer et al., 2018), genetic distance among samples and the reference may be an important factor to consider.

The detection of genomic polymorphisms is affected by the evolutionary time between the individuals being sequenced and the reference genome (Bohling, 2020; Prasad et al., 2022; B. N. Reid et al., 2021). This has implications on both the detection and the genotyping of single nucleotide polymorphisms (SNPs) and structural variants (SVs). For example, variant calling through pipelines such as GATK and freebayes use Bayesian inference to call genotypes (i.e., the likelihood of a genotype, given the data; Auwera et al., 2014; Garrison & Marth, 2012). When high differentiation between reference genome and the sampled individuals exists, a high proportion of segregating sites will emerge as fixed differences between the samples and the reference genome (i.e., homozygote non-reference in diploids), and therefore uncertain genotypes can have a higher likelihood of being called as homozygote non-reference, even if this is not the case. Analogously, most methods used to detect SVs (read-depth, split paired-read, breakpoints, and assembly) compare mapped reads to the reference genome (Pirooznia et al., 2015). Several studies on humans have demonstrated that ethnicity-specific reference genomes are beneficial (Ameur et al., 2017; Dewey et al., 2011; Fakhro et al., 2016; Lacaze et al., 2019). Specifically, the targeted reference genomes improved reliability of genetic and structural variation calls (Ameur et al., 2017; Fakhro et al., 2016). Moreover, recent studies have demonstrated that increasing phylogenetic distance between target species and reference genome decreases mapping efficiency and has strong effects on evolutionary inferences made from the data (Bohling, 2020; Prasad et al., 2022).

The three-spined stickleback (*Gasterosteus aculeatus*) is a supermodel in evolutionary biology (K. Reid et al., 2021), and research on this small teleost fish has pioneered discoveries related to the genomics of adaptation (Feulner et al., 2015; Haenel et al., 2019; F. C. Jones et al., 2012; Roesti et al., 2015), adaptive divergence (Feulner et al., 2015; Huang et al., 2016; Roesti et al., 2013) and development (Shapiro et al., 2004; Spitz et al., 2001). Genomic investigations using *G. aculeatus* have mostly relied on a single high-quality chromosome-level reference genome from an isolated population in Alaska (F. C. Jones et al., 2012), which has been updated with multiple improvements (Glazer et al., 2015; Nath et al., 2021; Peichel et al., 2017; Roesti et al., 2013). Recently, several additional *de novo G. aculeatus* contig-level genome assemblies have been made available, including European and marine derived assemblies (Berner et al., 2019). These additional genome assemblies may be appropriate given the wide geographic distribution of the species; there are Atlantic and Pacific clades of *G. aculeatus* that diverged an estimated 44.6 Kya with extensive phenotypic and genetic diversity across its range (Fang et al., 2018; McKinnon & Rundle, 2002).

In this study, we report the generation and *de novo* annotation of a European *G. aculeatus* genome assembly derived from a gynogen individual (Samonte-Padilla et al., 2011). The gynogenesis process in *G. aculeatus* produced a near complete homozygous diploid fish (Samonte-Padilla et al., 2011), which helps alleviate some of the genome assembly difficulties associated with heterozygosity. Then, using European and North American derived reference genomes, we investigated the effect of reference genome origin. We used high-quality genome-wide resequencing data from 60 *G. aculeatus* individuals from 5 recently diverged lake-river pairs distributed across the European (Atlantic clade; Fang et al., 2018) and North American (Western North America; Pacific clade) *G. aculeatus* ranges. The effect of using a local reference genome was investigated through comparing mapping efficiency (i.e., the percentage of reads that are successfully mapped to the reference genome), genotype calling, genome scans for *F*_ST_, Tajima’s *D*, and π, and identification of SVs. The resequenced genome data have been used to investigate the distribution of islands of differentiation across the genome (Feulner et al., 2015) and the role of copy-number variation in adaptation (Chain et al., 2014), offering baselines to evaluate specific metrics against a new local genome. We also compared how the different genomes affect DNA-methylation calling, focusing on an additional 50 European fish for which reduced-representation bisulfite sequencing (RRBS) was available (Sagonas et al., 2020). We hypothesise that the mapping and genotyping results will generally fall into 3 categories: (*h*_*1*_) a local reference genome has a significant and putatively benficial effect, (*h*_*2*_) one reference genome is better than the other, and (*h*_*3*_) a local reference genome has an effect, but the effects are more substantial with local populations (i.e., a mixture of both *h*_*1*_ and *h*_*2*_). Our hypotheses can be observed in the outcome of statistical tests: *h*_*1*_ presents as a significant interaction between reference origin and population origin, *h*_*2*_ as a significant effect of reference origin, and *h*_*3*_ presents similarly to *h*_*1*_ with a significant interaction, but the post-hoc analysis indicates a clear bias towards one of the reference genomes.

## Material & Methods

### Reference Genome Assembly and Annotation

The induction of the diploid gynogen individual followed the protocol established for three-spined sticklebacks (Samonte-Padilla et al., 2011). In brief, fertilised stickleback eggs were mixed with UV-irradiated sperm (2 minutes of exposure) and then exposed to a heat-shock treatment of 34°C for 4 minutes, 5 minutes post-fertilisation. The treatment caused the genetic inactivation of the sperm, resulting in homozygote maternal offspring that lack paternal alleles (Samonte-Padilla et al., 2011). In order to increase the likelihood of embryo development, two siblings from the same family were used for this process. After 5 months post hatching, a fish was sacrificed. DNA was extracted using the Qiagen high molecular weight extraction kit following the manufacturer’s protocol. Sequencing was then conducted at the Beijing Genomics Institute (BGI), taking place on a PacBio platform. In total, 2,805,993 reads were generated with a coverage of 44.1X. A total of 214,285 PacBio reads were discarded before further analysis given their short length. In addition, Illumina paired-end sequencing libraries in a HiSeq2500 platform were constructed with insert sizes of about 170 base pairs (bp), 500 bp and 800 bp.

Genome assembly was performed using Canu v1.6 assembler (Koren et al., 2017), followed by an internal polishing step using Quiver. To hybrid polish the PacBio assembly, a total of 316,797,342 high-quality Illumina reads were mapped to the contigs using BWA-MEM v0.7.15 (Li, 2013) and the alignment was then used for further polishing using Pilon v1.22 (Walker et al., 2014). Illumina raw reads were trimmed, and low quality and adaptor sequences were removed using Cutadapt v1.13 (Martin, 2011). To evaluate the PacBio *de novo* contig-level assembly and search for potential misassemblies, we used FRCbam v1.3.0 (Vezzi et al., 2012), whereas its completeness in terms of core orthologous genes was assessed using BUSCO v3.0.1 (Simão et al., 2015) and the ‘actinopterygii’ data set. The 32.1kb contig tig00001189_pilon mapped to the mitochondrial genome of the North American *G. aculeatus* (Peichel et al., 2017) and was trimmed and labelled the mitochondrial genome. The sequence was trimmed to only include the 15.8kb that aligned to the North American mitochondrial genome given the size of the mitochondrial genomes are moderately conserved even across long phylogenetic distances (Gissi et al., 2008). The excess 16.3kb of contig tig00001189_pilon were labelled and kept in the assembly, making up two additional contigs.

To scaffold the European *G. aculeatus* contig-level gynogen assembly into pseudo-chromosomes we used Chromosemble from the Satsuma2 package (Grabherr et al., 2010). Here, we ordered and oriented the contigs based on synteny with the North American *G. aculeatus* chromosome-level genome (Peichel et al., 2017), excluding the unmapped scaffolds. We tested for the effect of contig size on alignment rates, and only retained contigs with an alignment rate of 70% or above for further assembly. Gynogen contigs not scaffolded onto a pseudo-chromosome were concatenated in size order into an unmapped scaffold for population genomic analyses, separating each contig by 1,000 N’s. Synteny between the European and North American *G. aculeatus* assemblies was calculated using the SatsumaSynteny2 function (Grabherr et al., 2010), and plotted using the Circos v0.69 visualisation tool (Krzywinski et al., 2009). It should be noted that the reference genomes have been created using distinctly different methodologies. As such, significant reference origin by population origin interactions that do not clearly exhibit a reciprocal effect in the test statistic likely include a genome quality effect.

Repetitive sequences were identified *de novo* in the European pseudo-chromosome assembly using Repeat Modeler v.2.0.1 while Repeat Masker v.4.1.1 (Smit et al., 2015) was used to mask the genome using the three-spined stickleback and zebrafish libraries in two separately rounds. The results of each round were then analyzed together, complex repeats were separated, to produce the final repeat annotation. Genome annotation was performed on the European repeat-masked pseudo-chromosome genome assembly using MAKER2 v.2.31.9 (Holt & Yandell, 2011). Subsequent genome annotation was performed following a two-round approach. For the first round, the repeat annotation data (release 95) as well as *G. aculeatus* transcriptome and protein sequences from ENSEMBL and UniProt/SwissProt databases were used as evidence sets for the prediction of gene models, while *est2genome* and *protein2genome* setting were set as 1. For the second round, SNAP (Korf, 2004), with ADE of 0.25 and length of 50, and AUGUSTUS v.3.2.3 with default values (Stanke et al., 2008) were trained on the gene model predicted from the first round. Functional annotation was performed using BlastP against UniProt proteins with an E-value threshold of 1e-5, and InterProScan v.5.4-47 (P. Jones et al., 2014) was used for domain annotation. The resulting gene models were filtered to retain those with AED value of 0.5 or less, having PFAM annotations and significant hits to known proteins against UniProt DB (E-value 1e-5).

We identified orthologous and paralogous gene families among the North American and European *G. aculeatus* reference genomes using OrthoFinder v2.4.1 with the default parameters (Emms & Kelly, 2019). Protein sequences were extracted using the getfasta function in the BEDtools toolset v2.26.0, using the *-split* parameter to only include exonic regions (Quinlan & Hall, 2010). Where applicable, downstream analyses were restricted to only include the 7,529 1-to-1 orthologues identified between the two assemblies to remove any biases stemming from differences in gene number or functional annotation.

### SNP Data Collection and Processing

Whole genome resequencing data from a total of 60 *G. aculeatus* individuals were used from 5 recently diverged freshwater lake (_L) and river (_R) population pairs (table S1; details on sampling, library preparation, sequencing, and original data processing up to adapter trimming can be found in; Feulner et al., 2015). The five population pairs were sampled from two sites in Germany (G1 and G2; European; Atlantic clade), and one site in Norway (No; European; Atlantic clade), the United States (US; North American; Pacific clade), and Canada (Ca; North American; Pacific clade).

For the genome scan using SNP data, raw data was processed following previous procedures (Feulner et al., 2013). Adapter-cleaned reads were trimmed using Trimmomatic v0.36 (Bolger et al., 2014) in paired-end mode, trimming read tails with a PHRED quality score below 20 and trimming to a maximum of 50bp. Using BWA-MEM v0.7.17 (Li, 2013), reads were independently mapped to both the anchored European and US reference genomes (Peichel et al., 2017). Mapping efficiency was calculated using Bamtools v2.4.1 (Barnett et al., 2011). All downstream processing of mapped reads and methodology for variant calling were identical for both reference genomes. Mapped reads were processed using Picard toolkit v2.18.7 (https://broadinstitute.github.io/picard/), applying FixMateInformation and CleanSam. All reads belonging to the same individual from different lanes were combined using MergeSamFile, and then duplicate reads were flagged using MarkDuplicates. Variant calls were performed using GATK v4.0.6.1 (McKenna et al., 2010), calling variants in all genomes simultaneously, split by chromosome. The final set of SNPs was produced using hard filtering following the best practise workflow (see supplementary methods for filtering thresholds; (Depristo et al., 2011). Genome mapping and variant calls were conducted on the QMUL Apocrita High Performance Computing Cluster (King et al., 2017).

The gynogenesis process employed here aims to purge heterozygous variation, enabling us to reconstruct complex genomic regions. To assess the proportion of the genome that remains heterozygotic we used the same GATK variant calling and filtering pipeline detailed above with the paired-end Illumina libraries used to polish the gynogen assembly.

### Phylogenetic Analyses

In order to assess the phylogenetic relationship between the reference genomes and the 60 resequenced genomes, we compared maximum-likelihood phylogenies based on each variant call with its respective reference genome. Maximum-likelihood phylogenies for both SNP calls were inferred using RAxML v8.2.11 (Stamatakis, 2014). We randomly sampled 1% of all segregating sites by setting the “—select-random-fraction” parameter to 0.01 in the GATK function SelectVariants. The resulting VCFs were converted to PHYLIP format (Felsenstein, 1989) using the vcf2phylip.py script (doi:10.5281/zenodo.1257058). Trees were constructed using the GTRGAMMA model with 1000 bootstraps. Phylogenetic trees were plotted using the R package *ggtree* package v2.2.4 (Yu et al., 2017).

### Estimations of Genotype Bias

Calling of genotypes (i.e., heterozygote versus homozygote) may be affected by reference genome origin. Measures of genome-wide zygosity across all segregating sites were performed by parsing the genotype field in the VCF file using a custom R script. Genotypes were grouped into 5 distinct categories: homozygous reference, homozygous non-reference, heterozygous reference/non-reference, heterozygous non-reference/non-reference, and missing. This process was repeated to estimate genotypes for each individual mapped to both reference genomes.

### Genome Scan Using SNP Data

Genome scans were performed in R v4.0.2 (R Development Core Team, 2019), and population genetics indices were calculated using the R package *PopGenome* v2.7.5 (Pfeifer et al., 2014). We measured Tajima’s *D* and π in each population, and *F*_ST_ in each parapatric population pair. All metrics were calculated in non-overlapping 20kb windows across all 20 autosomal chromosomes and the unmapped scaffold. To obtain outlier genomic windows, we extracted the top 1% of the empirical distributions for each metric and population (or population pair for *F*_ST_). This conservative criterion (e.g., Feulner et al., 2015; Lai et al., 2019; Stern & Lee, 2020) was chosen to increase our confidence in defining outliers. Finally, we defined outlier genes as any gene overlapping one or more outlier genomic windows from the SNP-based genome scans using the *foverlaps* function in the R package *data*.*table* v1.9.6 (Dowle et al., 2015).

### Detection of Structural Variants

To investigate the impact of reference genome origin on the detection of structural variants (SVs), we used two independent SV callers, DELLY2 v0.8.3 (Rausch et al., 2012) and LUMPY v0.3.0 (Layer et al., 2014). Both LUMPY and DELLY were run using the default parameters. The LUMPY call was genotyped using SVTyper v0.7.1 (Chiang et al., 2015), and all SVs marked with the “LowQual” DELLY flag were removed. To ensure SVs found by both programs were the same, only SVs with an overlap of at least 50% were accepted and merged into a cohort level VCF file using SURVIVOR v1.0.3 (Jeffares et al., 2017). For downstream analyses, we included autosomal duplications, deletions, and inversions with a length of between 1kb and 1Mb supported by at least 6 split or discordant reads. All samples that were homozygote non-reference or heterozygote across all samples were analysed separately as these variants likely arise from the reference genome assemblies. Genes with coordinates entirely nested within SVs were defined as structurally variable genes (SVG) using the *foverlaps* function in the R package *data*.*table* v1.9.6.

### Methylation Data Processing and Genome Scan

For the DNA methylation analysis, we used 50 methylomes of laboratory full-sib families of European *G. aculeatus* obtained from the study of Sagonas et al. (2020), were we investigated whether parasite infection alters genome-wide patterns and levels of DNA methylations. For each fish, a single-end library of 100 bp reads with an average of 11.5 million reads was produced. The raw data were quality checked using FASTQC v0.11.5 (Andrews, 2010). Cutadapt v1.9.1 (Martin, 2011) was used to trim and filter low-quality bases (−q 20), remove trimmed reads shorter than 10 bases and remove adapters using multiple adapter sequences (ATCGGAAGAGCACAC, AGATCGGAAGAGCACAC and NNAGATCGGAAGAGCACAC) with a minimum overlap of 1 bp between adapter and read. Trimmed reads were independently mapped against the European and US reference genomes (Peichel et al., 2017) to extract methylated cytosines, using Bismark v0.22.1 (Krueger & Andrews, 2011) with the Bowtie2 v2.3.2 aligner, allowing up to 2 mismatches. Similar to the SNP data, Bamtools v2.4.1 (Barnett et al., 2011) was used to calculate the mapping efficiency.

Cytosine methylation ratios in CpG sites was estimated for each fish and differentially methylated sites (DMS) were calculated between the two treatment groups (no parasite exposed or exposed to the nematode parasite *Camallanus lacustris*) respectively (for more information, see Sagonas et al., 2020) using the R package *MethylKit* v1.14.2. CpG methylation ratios were estimated by calculating the number of reads mapping to a given position carrying a cytosine divided by those reads carrying either C or T. CpG sites with coverage below 10X and sites that have more than 99.9th percentile of coverage were discarded in each sample from downstream analyses. We selected DMS between treatments using the following criteria: change in fractional methylation larger than 15%; q-values lower than 0.01 (SLIM method), and presence of methylated Cs in at least 50% of the samples within a treatment group. Differentially methylated genes were identified using the *genomation* R package v.1.1.0 (Akalin et al., 2015); a gene was considered differentially methylated if at least one DMSs was located no further than 1.5-kb upstream and 500 bases downstream of it.

### GO Enrichment Analysis

To examine how the origin of the reference genome impacts functional enrichment of outlier genes, structurally variable genes, and differentially methylated genes, we performed gene ontology (GO) enrichment analyses. GO enrichment analyses were performed using the g:GOSt function in the g:Profiler v0.1.9 package (Raudvere et al., 2019). Outlier genes were grouped by all combinations of reference genome origin versus population of origin, resulting in 4 distinct combinations: Europe-Europe, Europe-North America, North America-Europe, and North America-North America. The lists of outlier genes or SVGs identified using the European assembly were assigned the orthologous North American gene identifiers for GO enrichment. *P*-values were corrected for multiple testing using FDR.

### Statistical Analyses

Linear mixed effect models in the *lme4* package (Bates et al., 2015) were used to analyse how the origin of the reference genomes affected mapping efficiency, detection of genome-wise zygosity, and population-genetic indices (i.e., Tajima’s *D*, π, and *F*_ST_). For mapping efficiency, the proportion of mapped reads that were singletons or duplications were independently used as response variables. An interaction between reference genome origin (i.e., North America or Europe) and population origin (i.e., North America or Europe) were assigned as fixed effects, and population ID was set as a random factor. The same approach was used for all analyses, independently replacing the response variable with the zygosity categories or population-genetic indices, retaining the same fixed and random effects. *P*-values were inferred using Satterwaite’s degree of freedom method in the *lmerTest* package (Kuznetsova et al., 2017). Tukey HSD post-hoc tests were performed using *emmeans* (Lenth et al., 2020).

## Results

### Reference Genome Assembly

The contig-level European gynogen assembly is 458.4Mb, making it 1.0% smaller than the North American reference genome (463.0Mb; Peichel et al., 2017). We achieved an N50 of 0.746Mb, comprised of 1,906 contigs (table 1). Guided by synteny, we conservatively placed 419.3Mb (91.5%) into 21 chromosomes (fig. 1), forming a pseudochromosome level assembly allowing for a more accurate comparison of the impact of the origin of reference genomes on population genomic metrics. A combination of gene evidence from the *G. aculeatus* North American reference genome (available cDNA and protein sequences) and *ab initio* gene predictions resulted in 22,739 genes annotated and a BUSCO completeness score of 95.2%. A total of 18,255 (90.2%) genes in the European gynogen assembly were orthologous to genes in the North American assembly. Additionally, we confirmed the European and North American derived reference genomes are nested within the phylogeny of populations sampled from the same geographic regions (supplementary fig. S1).

**Table 1.**
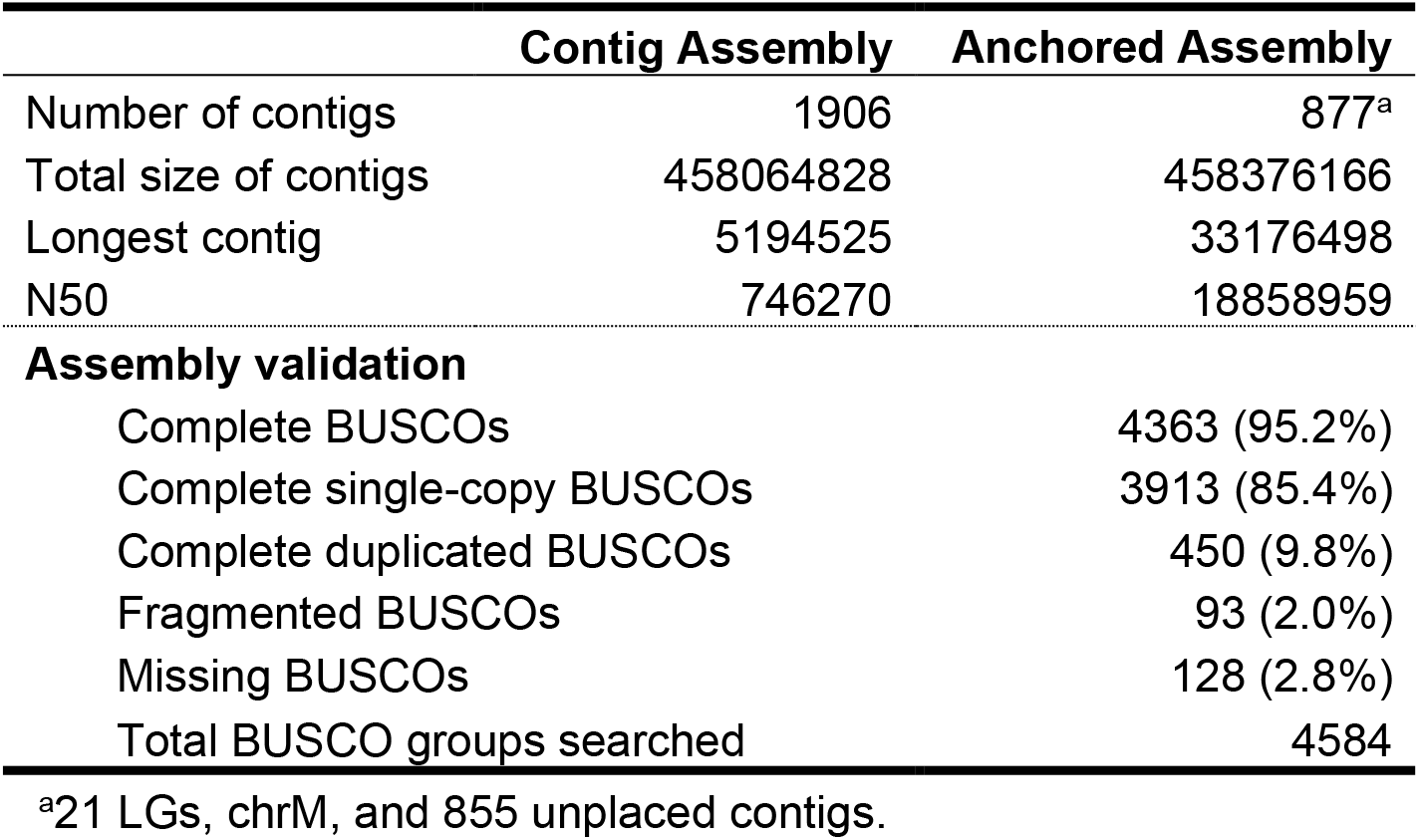
Assembly Statistics

**Figure 1.**
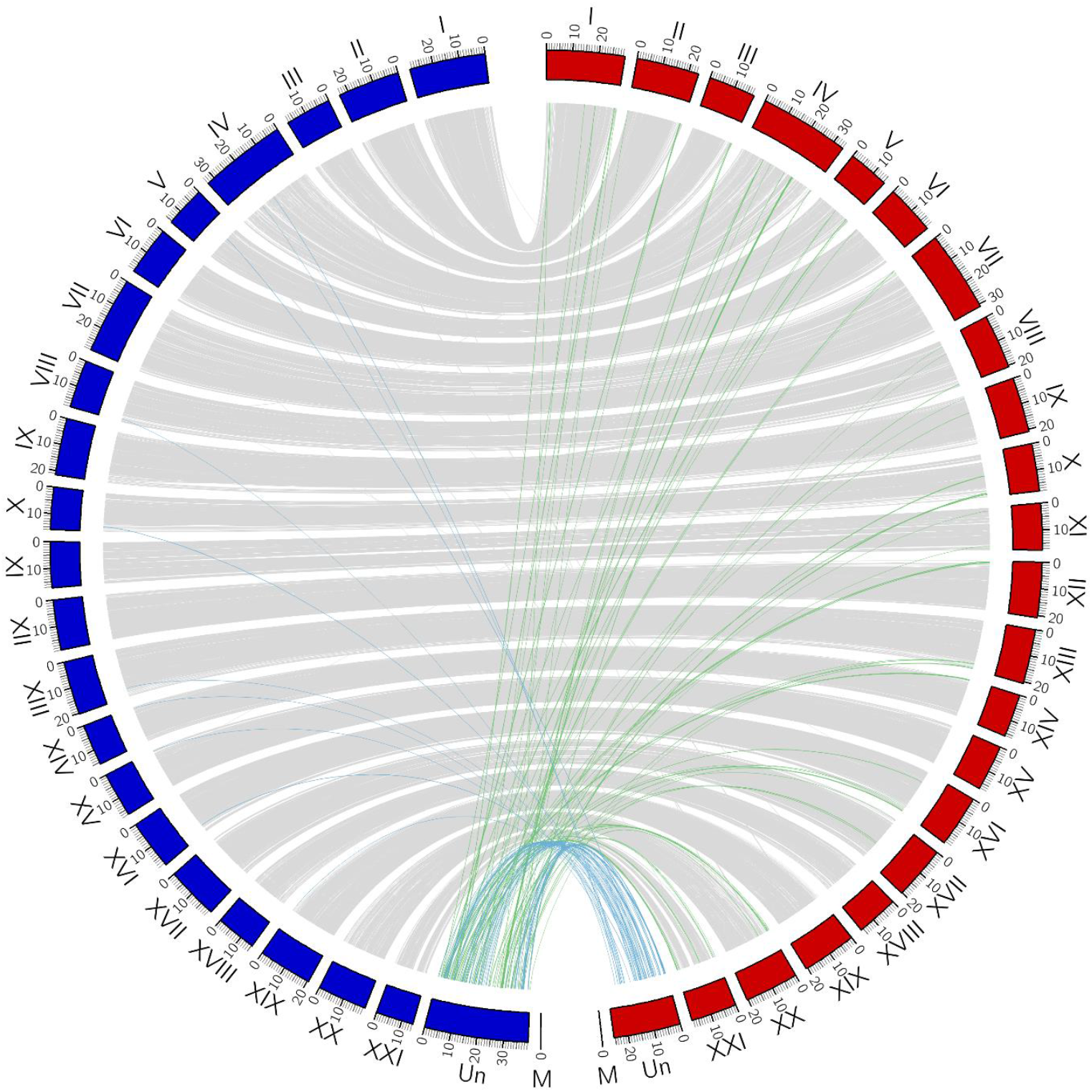
Synteny plot between the two *G. aculeatus* genome assemblies. Grey lines between the autosomes represent +99% syntenic blocks greater than 1kb between the European gynogen assembly (left, blue blocks) and the North American *G. aculeatus* assembly (right, red blocks). Coloured lines represent synteny between resolved regions and the unmapped scaffold in each assembly. Specifically, blue and green lines represent 1kb +99% syntenic blocks between the unmapped scaffolds and the alternate reference autosome.

Next, we called SNPs in the paired-end libraries used to polish the European genome assembly to assess the remaining heterozygosity in the gynogen genome assembly, which is expected to be homozygous after the gynogenesis procedure. In total we identified 299,931 SNPs (average genome-wide read depth of 60.6) in the genome: 3,118 SNPs were homozygote non-reference, and 296,813 were heterozygote. Hence, after two generations of gynogenesis only 0.07% of the genome remains polymorphic, compared to 0.53% ± 0.01% (mean ± SE) across all samples mapped to both reference assemblies (supplementary table S1). These SNPs likely stem from paralogous sequences that were not identified by the genome assembler or from errors in the SNP calling process.

### Mapping Efficiency

To assess the impact of reference genome origin on downstream analyses, we started by mapping whole genome resequencing reads from 60 individuals to the reference genomes. In total, we identified 10,672,162 SNPs using the North American reference genome, and 10,757,204 SNPs using the European reference genome (supplementary table S1). Overall, we achieved an average genome-wide depth of coverage of 20.40 (range; 11.09 - 35.46) using the North American reference and 20.99 (11.30 - 36.14; supplementary table S1) using the European reference. Reads mapped more efficiently (i.e., a higher proportion of reads mapped) to the European reference genome regardless of the population of origin with total reads mapping 2.31% ± 0.16% (European; estimate ± SE) and 1.13% ± 0.20% (North American) more efficiently, albeit with a greater increase in efficiency for local samples (LMER; Reference Origin:Population Origin, F_108_=21.38, P<0.001; fig. 2A and supplementary table S2). This same result was also observed when only considering properly matched paired-end reads (i.e., paired reads where both reads map in the correct orientation; 0×2 flag), where reads mapped more efficiently to the European reference, but there was a greater increase in efficiency for local populations (European, 3.14% ± 0.22%; North American, 1.36% ± 0.27%; Reference Origin:Population Origin, F_108_=25.94, P<0.001; fig. 2A). There were also 0.83% ± 0.08% (European) and 0.24% ± 0.09% (North American) fewer singletons (i.e., where only one of the paired-end reads mapped) when mapping European or North American populations to the European reference (LMER; Reference Origin:Population Origin, F_108_=25.10, P<0.001; fig. 2B). Reference genome or population origin had no effect on mapping of duplicate reads (LMER; Reference Origin:Population Origin, F_108_=0.01, P=0.942; Reference Origin, F_108_=0.12, P=0.729; Population Origin, F_8_=3.76, P=0.088; fig. 2B). Overall, using the European derived genome assembly noticeably improved mapping efficiency irrespective of the sample origins, although the effects were noticeably more efficient for European populations. Such results are consistent with hypothesis *h*_*3*_, which describes that a local reference genome is beneficial, but the effects are generally more substantial when using the European reference genome.

**Figure 2.**
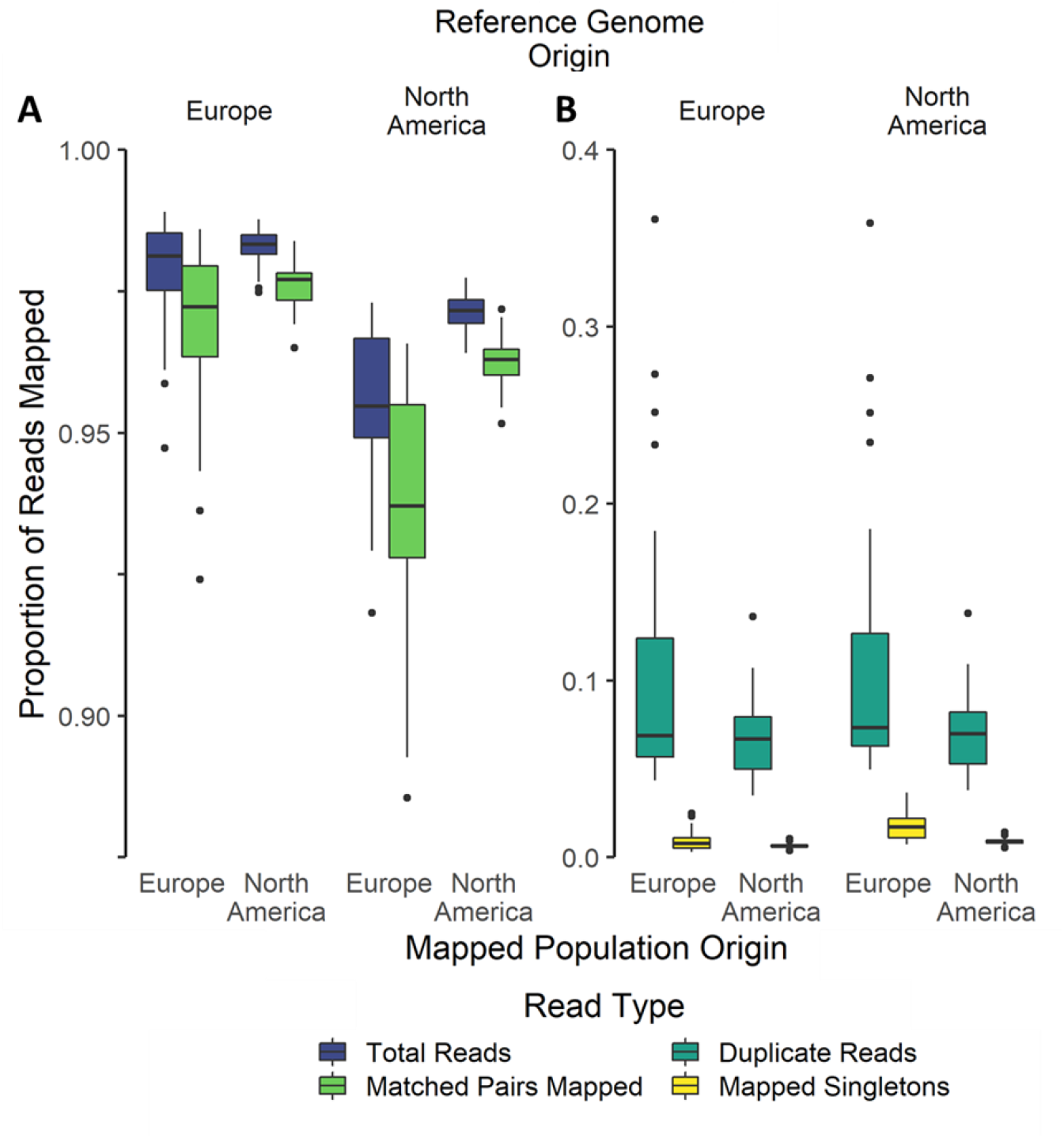
The effect of reference genome origin on mapping efficiency. Scales differ among panels, but the units are the same (*A*-*B*). There was significantly higher mapping efficiency when using the European reference genome regardless of sample origin for total reads (matched and singleton), matched pair reads (*A*) as well as significantly lower singletons (*B*). There was no difference in the number of duplicate reads identified. Reference genome origin is labelled at the top of the plot, and mapped population origin at the bottom.

### Estimations of Genotype Performance

When using a local reference genome, there were 6.24% ± 0.38% (European; estimate ± SE) and 0.47% ± 0.47% (North American) fewer missing genotypes (LMER; Reference Origin:Population Origin, F_428_=123.97, P<0.001; fig. 3 and supplementary table S2). The same was true for the detection of heterozygote genotypes of both classes (i.e., heterozygote reference/non-reference and non-reference/non-reference), where a local reference genome decreases the number of calls by 0.13% ± 0.09% (European) and 0.31% ± 0.11% (North American) for heterozygote reference/non-reference, and 0.10% ± 0.003% (European) and 0.05% ± 0.004% (North American) for heterozygote non-reference/non-reference genotypes (LMER; all models report the Reference Origin:Population Origin interaction; reference/non-reference, F_428_=9.86, P=0.002; non-reference/non-reference, F_428_=872.85, P<0.001; fig. 3). Conversely, we identified approximately 4 times higher proportions of homozygote reference genotypes when using a local reference genome for European populations (12.19% ± 0.37%) in comparison to the North American populations (3.36% ± 0.35%; LMER; Reference Origin:Population Origin, F_428_=691.30, P<0.001; fig. 3). Finally, approximately a two-fold decrease in the proportion of homozygote non-reference variants were identified when using a local reference genome for European populations (−5.72% ± 0.12%) over the North American populations (−2.53% ± 0.14%; LMER; Reference Origin:Population Origin, F_428_=2012.29, P<0.001; fig. 3). Except for duplications which were not significantly different among calls, using a local reference genome offers a benefit in the detection of every class of genotype. Overall, these results are consistent with hypothesis *h*_*1*_ which posited that a local reference genome would offer a benefit and would manifest as significant interactions.

**Figure 3.**
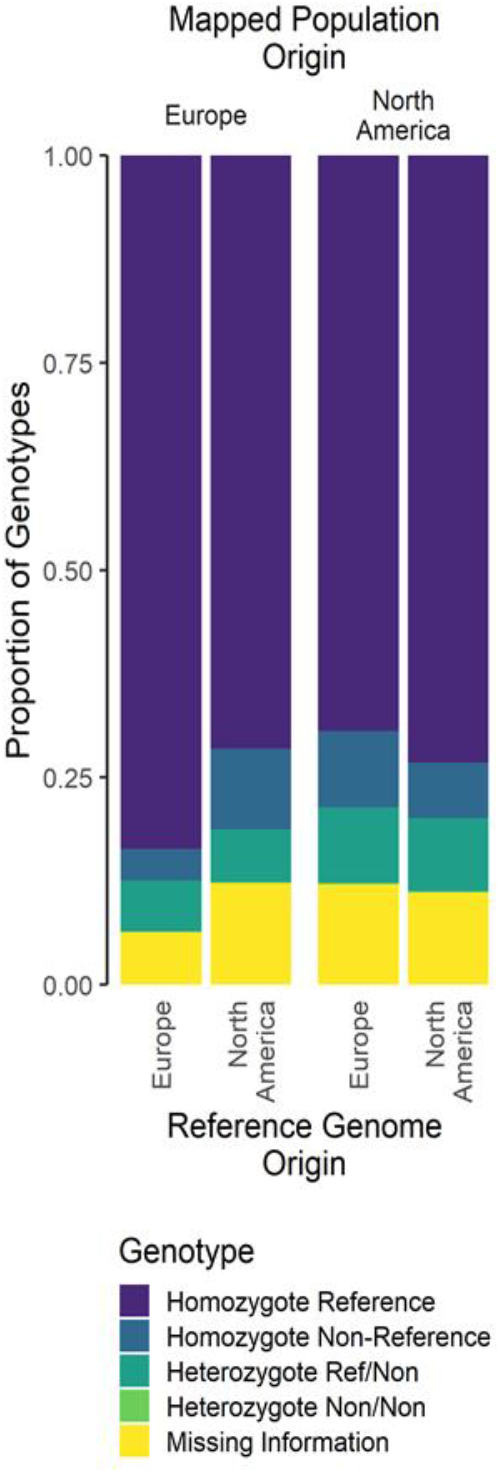
The proportion of SNP classes among all segregating sites. Segregating sites with no coverage or with a SNP that was removed during filtering are defined as missing information.

### Genome Scan Using SNP Data

The genome-wide distributions of metrics commonly used in population genomics were affected by the origin of the genome used (fig. 4 & S2-3). For a genome scan investigating patterns of differentiation, F_ST_ was 0.019 ± 8.4×10^−4^ (mean ± SE) higher for European populations and 0.005 ± 1.0×10^−5^ higher for North American populations when using a local reference genome (LMER; Reference Origin:Population Origin, F_220053_=324.60, P<0.001; supplementary table S2). A similar pattern was observed using T_D_, whereby T_D_ was 0.026 ± 2.95×10^−3^ (European) and 5.90×10^−3^ ± 3.61×10^−3^ (North American) higher when using a local reference genome (LMER; Reference Origin:Population Origin, F_406709_=63.97, P<0.001). In addition, using a local reference genome led to significantly lower values of nucleotide diversity being detected (LMER; Reference Origin:Population Origin, F_407698_=123.20, P<0.001), with low estimates for both European (−3.28×10^−5^ ± 5.39×10^−6^) and North American (−9.99×10^−5^ ± 6.60×10^−6^) populations. These results are consistent with hypothesis *h*_*1*_, where a local reference genome has a significant impact on analyses.

**Figure 4.**
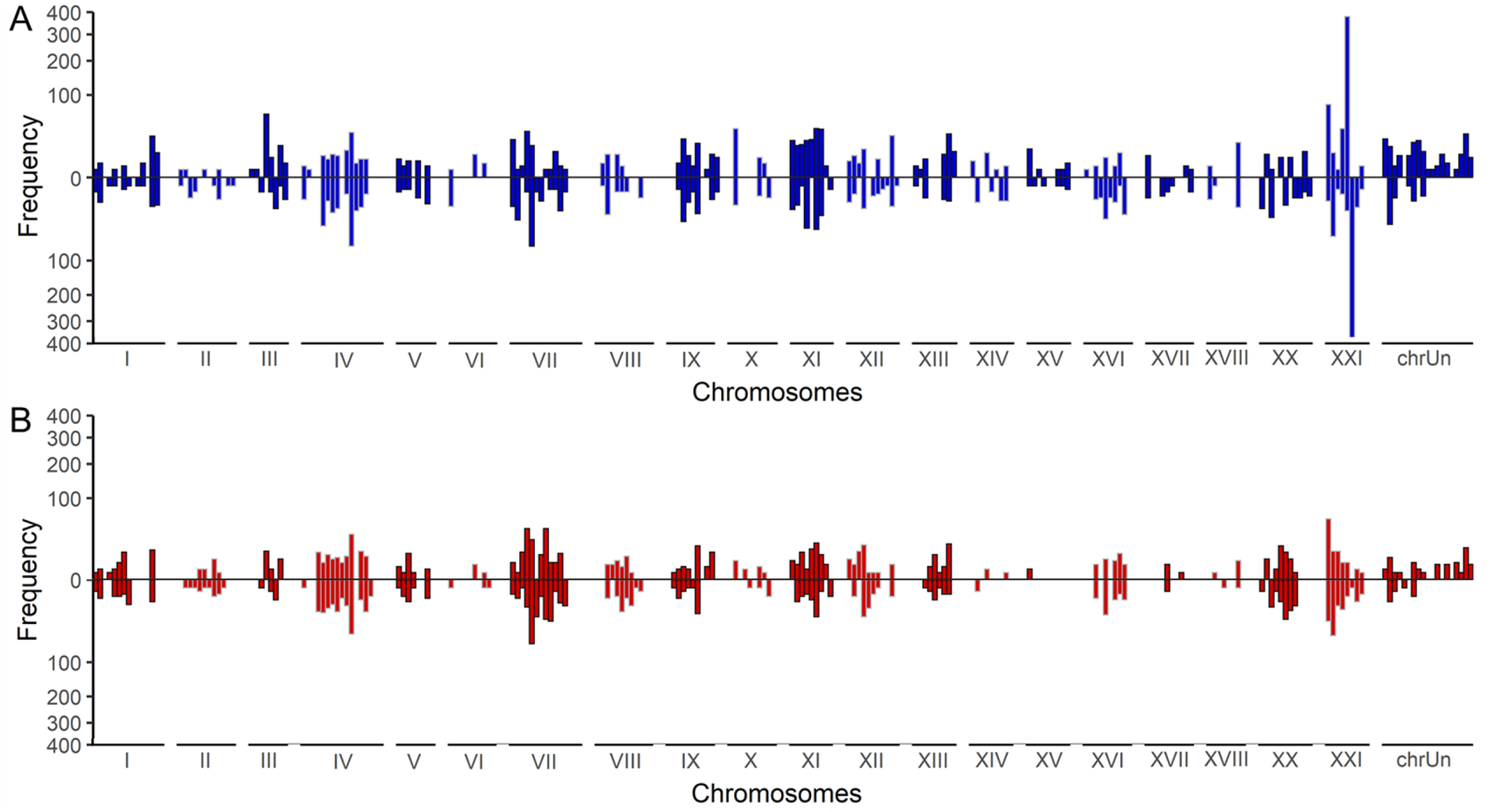
Comparing distributions of π outliers. Windows are compared across the genome for (top) European and (bottom) North American populations mapped to the (*A*) European or (*B*) North American reference genome. Axes are square root transformed.

Through sampling each metric individually and taking the top ∼1% of the F_ST_, π, or T_D_ distributions we generated multiple lists of outlier genes (table 2; supplementary table S3). Notably, the distributions of outlier genes across the genome were not significantly different when using either reference genome (two-sample Kolmogorov–Smirnov test; F_ST_, D=0.140, P=0.988; π, D=0.121, P=0.998; T_D_, D=0.157, P=0.962; fig. 4 & S2-3). Overall, higher number of outlier genes, including the subset of outlier 1-to-1 orthologous genes, were identified when using a local reference genome (table 2). The difference in genes among calls with different reference genomes putatively translated into no overlapping significantly enriched GO terms for the *F*_ST_, π, and T_D_ analyses if any enrichment was detected (table 2). Complete GO enrichment tables are reported in supplementary table S4.

**Table 2.**
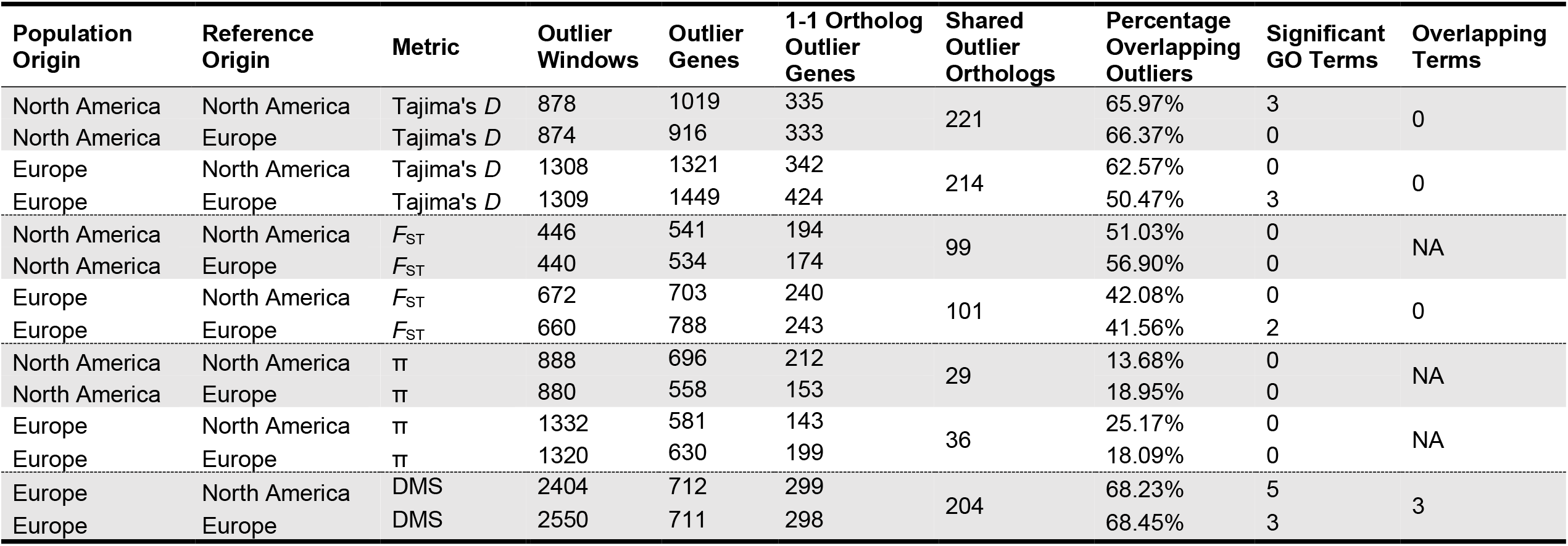
Distribution of outlier windows and differentially methylated sites (DMS), overlapping genes, and their functional enrichment.

### Genomic Structural Variants

We next addressed the question whether using a local reference genome affected the detection of structural variants. Firstly, we generated multiple distributions of SVs organised by the combination of reference genome and population origin (table 3). Overall, no fixed SVs were observed in all samples. The distribution of deletions significantly differed among SV calls (two-sample Kolmogorov–Smirnov test, D=0.278, P=0.008; fig. S4), whereby fewer SVs were detected when using the North American reference genome. The distribution of duplications and inversions did not significantly differ among SV calls with either reference genome (two-sample Kolmogorov–Smirnov test; duplications, D=0.109, P=0.890; inversions, D=0.140, P=0.754; fig. S4). Additionally, we identified less deletions when using a local reference genome, more deletions were identified overall when using the European reference genome (LMER; Reference Origin:Population Origin, F_108_=61.813, P<0.001; table 3, supplementary table S5-S6). Analogously, we identified significantly fewer inversions when using a local reference genome, but more inversions were identified overall when using the North American reference genome (LMER; Reference Origin:Population Origin, F_108_=34.52, P<0.001; table 3). Finally, a similar number of duplications were identified in European populations irrespective of reference origin, whereas fewer duplications were identified in North American populations when using a local reference genome (LMER, Reference Origin:Population Origin, F_108_=3.99, P=0.048; supplementary table S5-S6). Overall, the results of the SV analysis are in line with hypothesis *h*_*3*_ where the effects of a local reference genome are present, but unequal effects indicate one reference is better than the other.

**Table 3.**
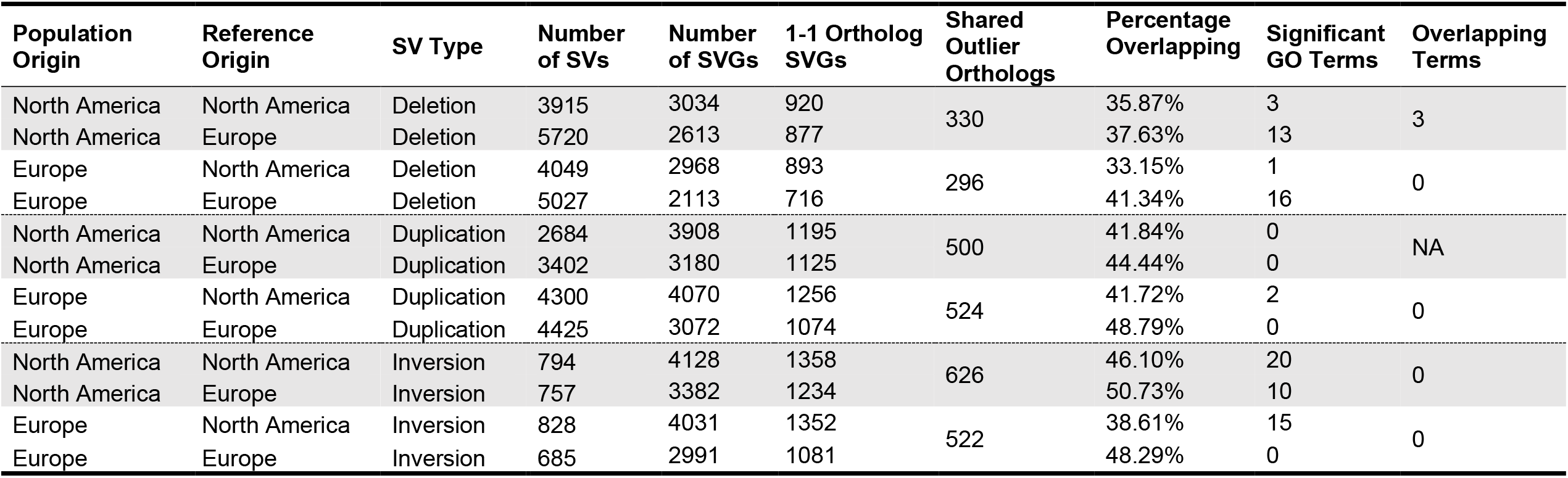
Distribution of structural variants and structurally variable genes and their functional enrichment.

| Population Origin | Reference Origin | SV Type | Number of SVs | Number of SVGs | 1-1 Ortholog SVGs | Shared Outlier Orthologs | Percentage Overlapping | Significant GO Terms | Overlapping Terms |
| --- | --- | --- | --- | --- | --- | --- | --- | --- | --- |
| North America | North America | Deletion | 3915 | 3034 | 920 | 330 | 35.87% | 3 | 3 |
| North America | Europe | Deletion | 5720 | 2613 | 877 |  | 37.63% | 13 |  |
| Europe | North America | Deletion | 4049 | 2968 | 893 | 296 | 33.15% | 1 | 0 |
| Europe | Europe | Deletion | 5027 | 2113 | 716 |  | 41.34% | 16 |  |
| North America | North America | Duplication | 2684 | 3908 | 1195 | 500 | 41.84% | 0 | NA |
| North America | Europe | Duplication | 3402 | 3180 | 1125 |  | 44.44% | 0 |  |
| Europe | North America | Duplication | 4300 | 4070 | 1256 | 524 | 41.72% | 2 | 0 |
| Europe | Europe | Duplication | 4425 | 3072 | 1074 |  | 48.79% | 0 |  |
| North America | North America | Inversion | 794 | 4128 | 1358 | 626 | 46.10% | 20 | 0 |
| North America | Europe | Inversion | 757 | 3382 | 1234 |  | 50.73% | 10 |  |
| Europe | North America | Inversion | 828 | 4031 | 1352 | 522 | 38.61% | 15 | 0 |
| Europe | Europe | Inversion | 685 | 2991 | 1081 |  | 48.29% | 0 |  |

Next, we investigated the role of a local reference genome on the detection of genes entirely nested within SVs, which we defined as structurally variable genes (SVG; table 3). This analysis was limited to 1-to-1 orthologs to ensure there was no bias arising from copy number variation among reference genome annotations. Here, we identified significantly fewer SVG-deletions for European populations using a local reference genome, however there was no significant difference in SVG-deletions in the North American populations when using either reference (LMER, Reference Origin:Population Origin, F_108_=25.94, P<0.001; supplementary table S5-S6). On the other hand, we identified significantly more SVG-duplications when using the North American reference, regardless of the origin of the population (LMER, Reference Origin:Population Origin, F_108_=9.40, P=0.003; supplementary table S5-S6). Finally, significantly fewer inversions were detected when using a local reference genome (LMER; Reference Origin:Population Origin, F_108_=62.55, P<0.001; supplementary table S5-S6).

To identify whether a local reference genome correlated with the detection of functional enrichment in SVs, we investigated the SVGs GO enrichment. Similar to the overall SVG distribution analysis, this analysis was restricted to 1-to-1 orthologs. We identified no overlap in functional enrichment in the majority of the comparisons (table 3). The one exception was SVG-deletions in the North American populations, where 3 out of 13 significantly enriched terms (signaling receptor regulator activity, signaling receptor activator activity, and receptor ligand activity) were identified in both calls. Overall, the detection of SVs and SVGs were affected in different ways by the origin of the reference genome.

### DNA Methylation Analysis

Finally, we conducted a DNA methylation analysis after mapping bisulfite sequencing reads to the two reference genomes. The inclusion of this analysis permits us to investigate the effect of reference genome origin on both DNA methylation analyses and on marker-based analyses, as opposed to the window-based genome scans reported above. The alignment of reads to the references showed significantly higher efficiency when using a local European reference genome (72.5%) compared to the North American reference (68.4%), resulting in an increase of 6% (paired t-test; t=-44.44, df=49, P<0.001). The more efficient mapping to a local reference produced a significantly higher calling of cytosine bases (6% increase, paired t-test; t=-28.31, df=49, P<0.001) and methylated Cs (8.2% increase, paired t-test; t=-33.18, df=49, P<0.001). Similarly, the comparison of the number of methylated sites per fish after filtering for low coverage sites revealed that using the local European reference genome resulted in identifying more methylated sites (560629.08 ± 12167.89) than with the North America reference (510240.14 ± 11037.78, paired t-test; t=-44.20, df=49, P<0.001). Additionally, the number of Differentially Methylated Sites (DMS) was higher when using a local European reference genome (N=2550 DMS) in comparison to the North American reference (N=2404 DMS). DMS overlapped with 711 and 712 genes in the European and North American assemblies, respectively, and specifically 298 and 299 genes were 1-to-1 orthologs between the two assemblies. A total of 204 of those genes with DMS (68%) were shared in both genomes. There were three significantly enriched GO terms (protein binding, calcium ion binding, and binding) shared in both analyses, and two enriched GO terms (cation binding and metal ion binding) only identified when using a divergent reference genome (Table 2).

## Discussion

Until genome-graphs are more widely available, individual reference genomes remain an integral part of population genomic analyses. However, the reference genome can introduce mapping biases that significantly influence downstream analyses and inferences (Bohling, 2020; Prasad et al., 2022; Valiente-Mullor et al., 2021). Meanwhile, the effects of using a local or more differentiated reference genome remains understudied for ecological and evolutionary model species. To address this knowledge gap, we generated a *de novo* annotated synteny-guided assembly of a European *Gasterosteus aculeatus* fish. Using this novel genome and the established North American reference genome to map sequence data of samples from different populations in Europe and North America, we confirm the reference genome origin significantly impacts downstream analyses. Most notably, a local reference genome increased mapping and genotyping performance. Specifically, mapping efficiency was significantly better using the European reference genome, but the increase in performance was greater for local samples. When using a local reference, more genomic sites were genotyped, and genome window-based estimates of T_D_ and *F*_ST_ increased whilst π slightly decreased. Similarly, structural variants (SVs) analysis gave slightly different results based on the reference genome used. Consequently, most GO analyses resulted in only a minor proportion of matching enriched GO functions when using different reference genomes. In contrast to the window-based methods, the marker-based DNA methylation analysis pipeline was relatively less affected by reference genome origin, but still about one third of differentially methylated genes was uniquely identified by one but not both references.

### Genome Assembly of a Gynogenetic Individual

Recent tools and techniques have increased the efficacy of reference genome assembly, such as utilising long and short read sequencing (Rhie et al., 2021), scaffolding with Hi-C (Peichel et al., 2017), optical or linkage mapping (Glazer et al., 2015), among the growing list of novel and effective techniques (Rhie et al., 2021). The generation of gynogenetic individuals purges genome-wide variation simplifying assembly of genomes (Christensen et al., 2018; Samonte-Padilla et al., 2011). Here, after applying a previously established protocol for gynogenesis, we putatively removed more than 99.9% of genome-wide variation, likely aiding in scaffolding by long-read sequencing. The contiguity of the resulting contig-level gynogen *G. aculeatus* assembly is comparable to the top few assemblies published by the Fish10k project (Fan et al., 2020) in terms of number of contigs and contig N50. Notably, there are a few altered placements of contigs into chromosomes among assemblies. The largest was the 2.07Mb contig tig00002041_pilon, which aligns to both the middle of chrVIII (9.25Mb-10.87Mb) and the end of chrIX (17.89Mb-18.29Mb) and was placed in Gy_chrIX by Chromosemble. The Atlantic and Pacific *G. aculeatus* clades diverged an estimated 44.6Kya (Fang et al., 2018) leaving substantial time for large genomic rearrangements to occur. As such the correct placement of tig00002041_pilon is similarly likely in either chromosome which illustrates the need for further investigations to resolve the differences among assemblies. Overall, the new European gynogen reference genome is high quality, enabling us to test the effects of reference genome origin on downstream population genomic analyses.

### Mapping and Calling Variants

Despite continuing advances in tools to assemble reference genomes and map sequenced reads, difficulty remains in correctly mapping reads to complex genomic regions enriched in heterozygosity, structural variation, or repetitive elements (Kajitani et al., 2014; Treangen & Salzberg, 2012). By using a reference genome with a longer evolutionary time to the most recent common ancestor (MRCA), complex variants have time to accumulate, likely decreasing mapping efficiency to genomic regions with arguably some of the most sought-after features (e.g., polymorphic regions associated with rapid evolutionary changes and adaptations). Here, we show that using a European stickleback reference genome that has a lower time to the MRCA with European populations increases mapping efficiency and decreases missing data. Conversely, the North American populations also showed increased efficiency when mapped to the European reference, but the difference was noticeably smaller than for European populations. Such a result indicates the removal of heterozygosity through gynogenesis (Samonte-Padilla et al., 2011) and the putative resolution of complex genomic regions in the European assembly improved mapping efficiency. These results are concordant with ethnicity-specific reference genome studies, which have demonstrated that local reference genomes increase depth of coverage resulting in increased sensitivity in variant calling (Ameur et al., 2017; Dewey et al., 2011; Fakhro et al., 2016).

### Genome Scans

The effects of improved mapping efficiency and a decrease in missing data were observed to have significant impacts on the estimation of important population genomic metrics. Here, genome-wide estimates of *F*_ST_ and T_D_ were higher when using a local reference genome, whereas the opposite pattern was true for estimates of nucleotide diversity (π). Despite the differences in genome-wide estimates, the genome-wide distribution of outlier windows of π, *F*_ST_, and T_D_ did not significantly differ across the genome for the same resequenced populations mapped to different reference genomes. Crucially, however, the detection of outlier genes was strongly impacted when using different reference genomes, even when conservatively limiting the analysis to 1-to-1 orthologs identified among the references. Specifically, genes overlapping π outliers had a low proportion of matching orthologs in the same populations when mapped to different reference genomes. Genes overlapping outliers from scans for *F*_ST_ and T_D_ were more consistent, but still revealed a large number of differences. In total, only 5.45% of the orthologs among assemblies were mapped to different chromosomes, clearly affecting the detection of outlier genes but only explaining a small proportion of differences observed. The difference in gene placement may instead highlight numerous resolved differences among assemblies that have accumulated since the divergence of Atlantic and Pacific *G. aculeatus* clades 44.6 Kya (Fang et al., 2018). For example, a small deletion in one reference assembly and not the other can result in a gene overlapping an outlier genomic window in only one scan. Overall improved mapping may help specific population genomic analyses including genome-wide representation sequencing where population structure may end up obscuring some SNPs present in only one or few populations (e.g., Baltazar-Soares et al., 2020).

### Structural Variant Detection

Similar to the effect of reference genome origin on SNP-based scans, the detection of SVs appears to be affected by both reference genome origin and the assembly regardless of origin. Firstly, fewer deletions and inversions were identified when using a local reference genome. This result follows expectations, as there is less time between MRCA for SVs to build up between the sampled population and the reference genome. Secondly, more deletions and duplications and fewer inversions were detected when using the European reference, irrespective of population origin, suggesting differences in the genome assembly plays a role in the detection of SVs. However, the differences in SV detection did not translate into any significant differences in the number of genes overlapping deletions. Overall, our results suggest the detection of SVs may be affected by a combination of mapping efficiency, time to MCRA, and the methods used to assemble a reference genome.

### Methylation

The most consistent analysis in terms of overlap among reference genomes was the marker-based DNA methylation test. Firstly, using a local reference genome significantly increased mapping efficiency, which resulted in more methylated sites and DMS being detected. The number of genes overlapping the DMS using either reference genomes was the most consistent among all analyses, with one fewer gene (0.14% of total genes) and ortholog (0.34% of total 1-to-1 orthologs) being identified when using a local reference. The DMS analyses recovered a relatively high proportion of overlapping outlier orthologs among reference genomes (68% of 1-to-1 orthologs), but still revealed an effect of the choice of reference. The higher proportion of overlap compared to window-based analyses may be due to the marker-based approach, which is less sensitive to genomic translocation, genome evolution, or assembly errors.

### GO Enrichment

The largest effect of reference genome origin was in the GO enrichment analyses of outlier genes from genome scans, with only a minor proportion of enriched GO terms overlapping when using different reference genomes. Given the overall small proportion of overlapping outlier genes, these results were to be expected. GO enrichment analyses are particularly sensitive to minor changes in the number of genes or annotations in lists of genes (Gaudet & Dessimoz, 2017). It should be noted, however, that to allow for a direct comparison of the effects of the different reference genomes, we focused on 1-to-1 orthologs. GO enrichment analyses are common in population genomics (e.g., Chain et al., 2014; Feulner et al., 2015; Reimegård et al., 2017; Liu et al., 2018), and our results highlight reference genome origin strongly impacts such inferences.

## Conclusions

Assembling reference genomes is a fast-moving field of research, which sees persistent updates and novel methodologies adopted (Rhie et al., 2021). Hence, variation seen in new reference genomes can reflect both the geographical range of the species distribution but also variation in methodologies used to sequence and assemble the genomes. For example, the two *G. aculeatus* genomes used here originate from samples representing two distinct lineages, the European and North American lineage but also differ significantly in how they were generated. Notably, the North American reference genome (Peichel et al., 2017) is part of a series of updates to the original *G. aculeatus* genome assembly (F. C. Jones et al., 2012). The original assembly used entirely Sanger sequence data (F. C. Jones et al., 2012), compared to the PacBio and Illumina sequence data used for the gynogen genome. As such, only when there is a significant interaction between population and reference origin and no obvious bias in the observations towards either reference can we exclude that the differences in sequencing and assembly methodologies are not the primary cause for the observed patterns.

The aim of this study was to investigate the effects of a local reference genome and its effects on downstream analyses. The reference-specific patterns of our results highlight that there is no simple solution. We suggest that the quality of the reference genome and annotations remains the single most important factor when choosing which reference to use. However, when multiple similar quality references are available, a local reference genome offers higher mapping efficiencies and decreases the proportion of missing data. The smallest reference effect among our analyses was for the marker-based methylation analysis, which had markedly more overlap among outliers in comparison to the window-based approach or the SV analysis, but still had over 30% of outliers that did not overlap. Taken together, using a local reference genome should increase the confidence of inferences made within a study, even if the difference is only minor.

## Data Accessibility and Benefit-Sharing

The raw genomic sequences of the 60 *G. aculeatus* fish were obtained from previous publications (Chain et al., 2014; Feulner et al., 2015), and retrieved from the European Nucleotide Archive, accession number ERP004574. The raw methylome sequences were obtained from a previous publication (Sagonas et al., 2020), retrieved from the NIH genetic sequence database, accession number PRJNA605637.

The European-derived gynogen reference genome assembly and their annotations are available from the European Nucleotide Archive (ENA) accession number PRJEB54679. Please note, to comply with ENA submission guidelines the unmapped chromosome (chrUn) was broken into individual contigs, and annotations were lifted over using a custom R script. A total of 316 gene annotations spanned contig boundaries in chrUn and were removed. The original chrUn sequence and annotations are available upon request.

Scripts used in the data preparation and analysis pipelines are available online (https://github.com/dthorburn/Origin_Matters).

## Acknowledgements

We thank Katie Peichel and Richard Nichols for their useful comments and discussion. We also thank Queen Mary University of London ITS Research support for technical assistance.

## Author Contributions

D.M.J.T. and C.E. conceived and designed the study. F.J.J.C., P.G.D.F., C.E., T.L.L., M.B.P., and I.E.S.P. obtained the samples and oversaw the sequencing. T.B.H.R., E.B.B. and M.M. created the Big Screen Consortium with grant support from the Max Planck Society to M.M. K.S. and M.B.P. oversaw the initial genome assembly and annotation. D.M.J.T. analysed the data, advised by C.E. D.M.J.T. and C.E. drafted the manuscript. All co-authors contributed to the final version of the manuscript. C.E. acknowledges the support of the German Science Foundation (DFG, EI841/4-1 and EI 841/6-1). D.M.J.T. was supported by a studentship from Queen Mary University of London.

