## Supplementary Figures for "Origin Matters: Using a Local Reference Genome Improves Measures in Population Genomics"

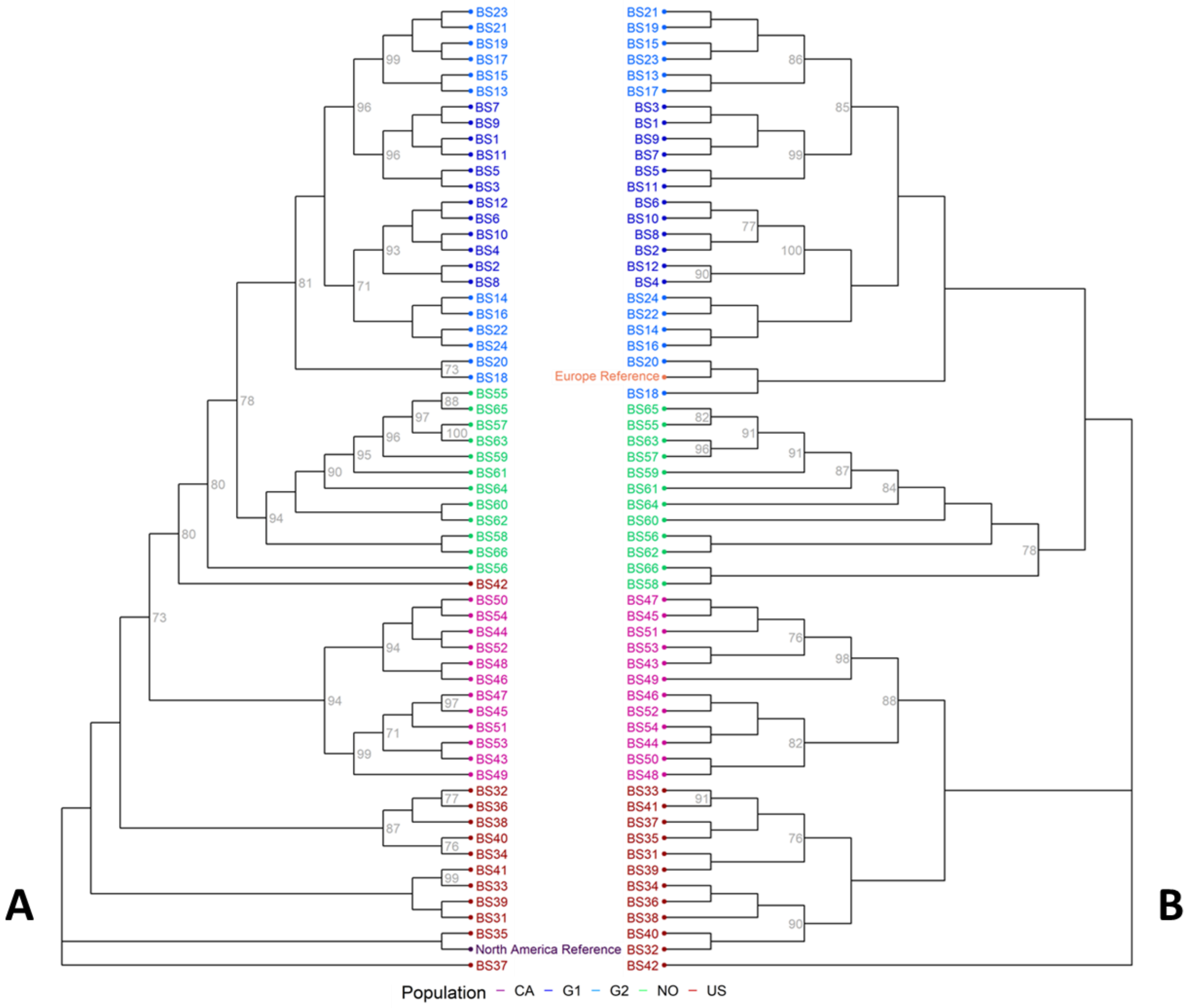


**Supplementary Figure S1.** Maximum likelihood tree based on polymorphism data called from [A] the North American reference genome and [B] the European reference genome. Both reference genomes were added to the phylogeny by randomly sampling 1% (~100k) of segregating sites across the genome. Nodes with higher than 70% bootstrap support are shown in grey at the node. Individuals are coloured by population-pair

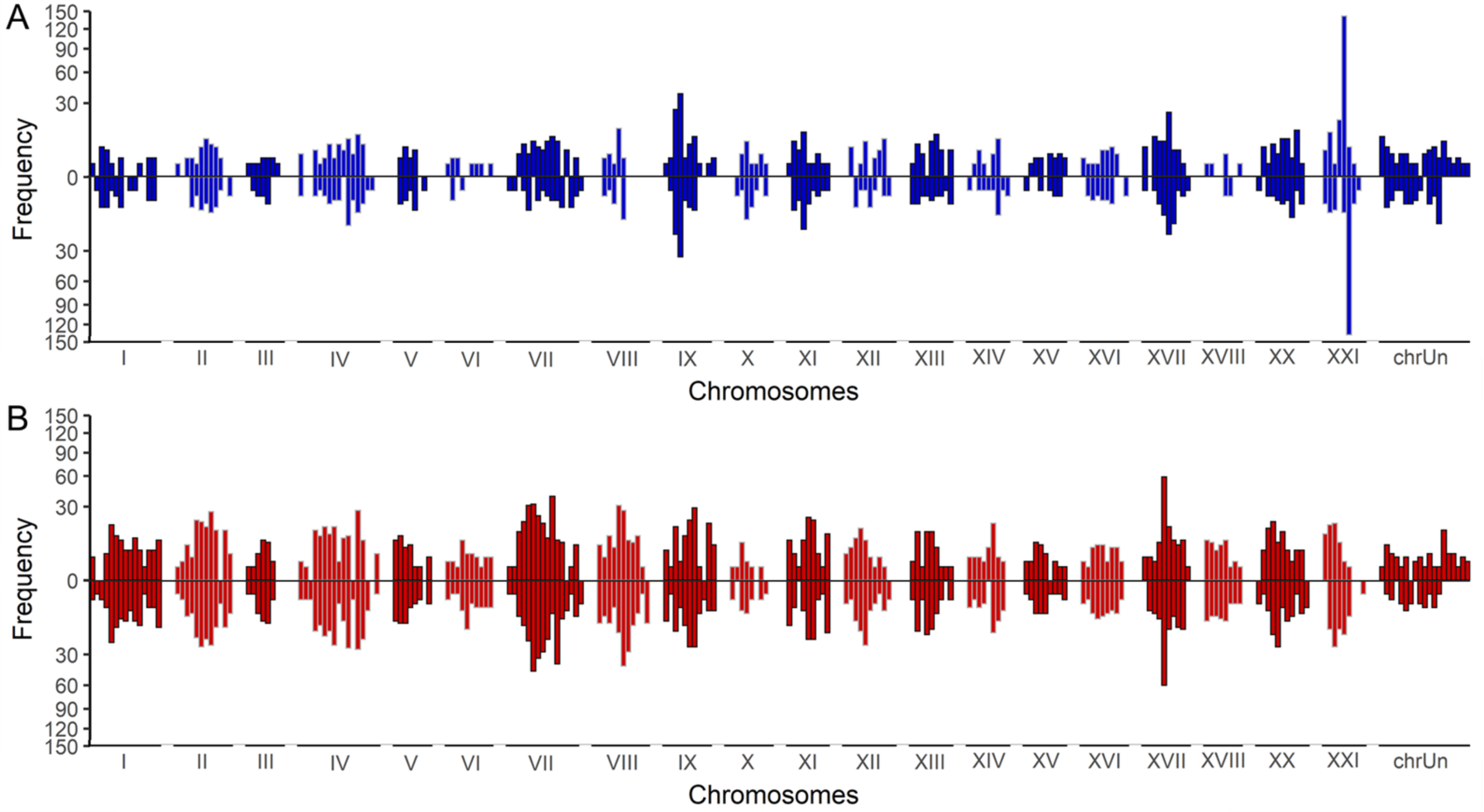


**Supplementary Figure S2.** Comparing distributions of outlier Tajima's *D* windows across the genome for (top) European and (bottom) North American populations mapped to the (*A*) European or (*B*) North American reference genome. Axes are square root transformed.

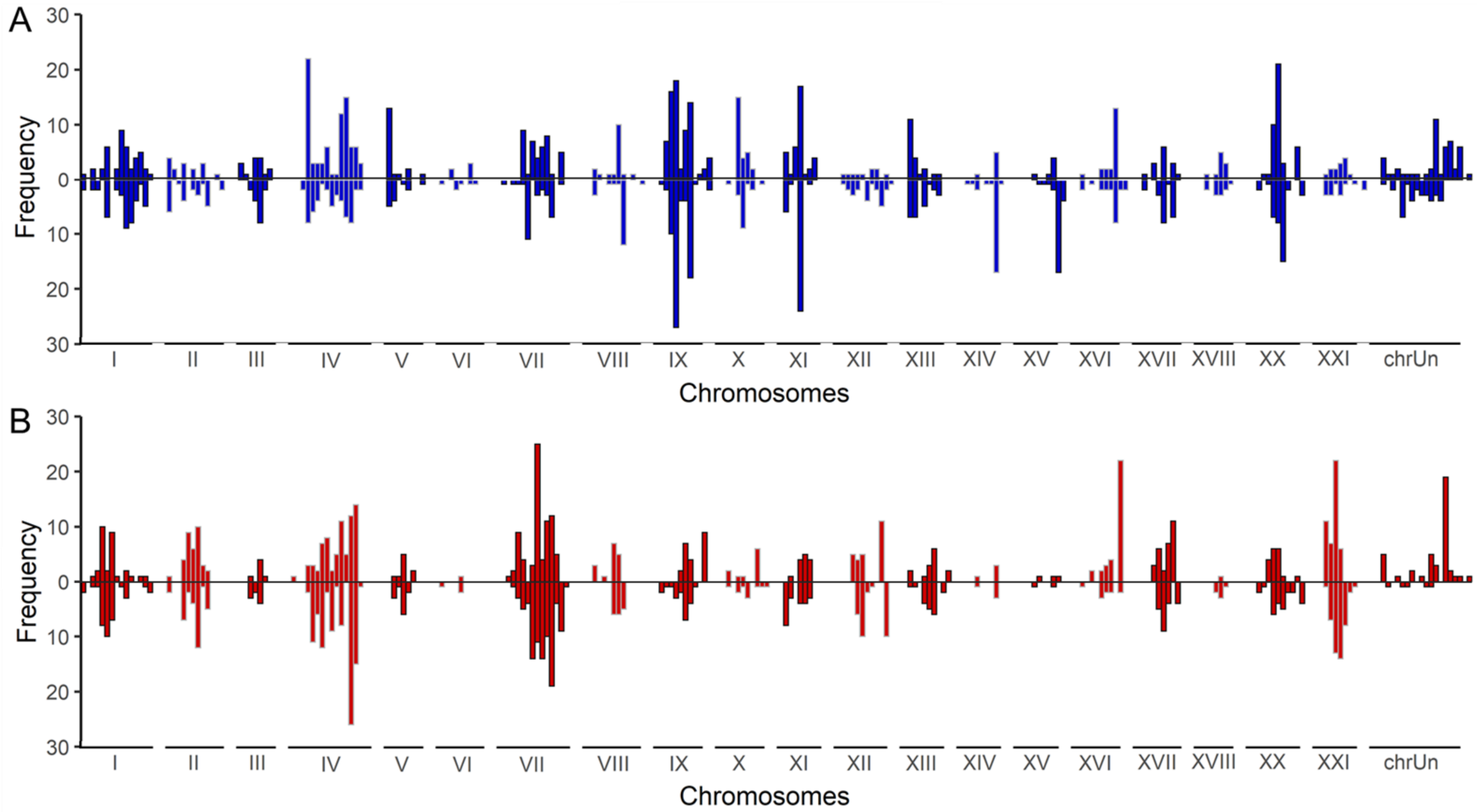


**Supplementary Figure S3.** Comparing distributions of outlier $F_{ST}$ windows across the genome for (top) European and (bottom) North American populations mapped to the (*A*) European or (*B*) North American reference genome. Axes are square root transformed.

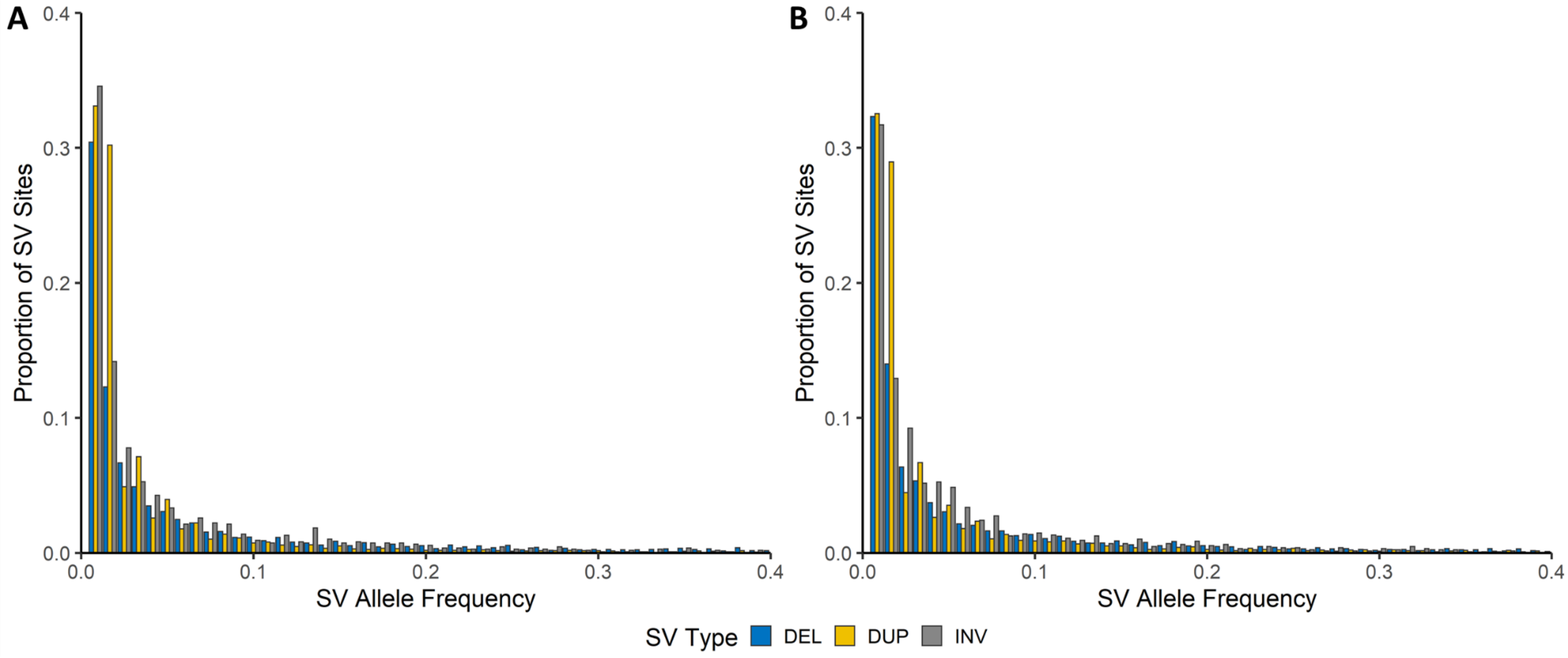


**Supplementary Figure S4.** The allele frequency spectrum of bi-allelic SVs across all 60 individuals, exhibiting that the majority of deletions (blue), duplications (yellow) and inversions (grey) occur at low frequencies when using both the (*A*) European or the (*B*) North American reference genomes.
